# Evaluation of BOLD effects in the rat cortex

**DOI:** 10.1101/2021.05.12.443862

**Authors:** Nathalie Just

**Author notes:** Corresponding author: Nathalie Just, PhD, HDR.

## Abstract

**Purpose:** This study aimed to characterize Blood oxygen level-dependent (BOLD) effects in ^1^H-MR spectra obtained during optogenetic activation of the rat forelimb cortex for the correction and estimation of accurate metabolite concentration changes.

**Methods:** T_2_*-induced effects were characterized by linewidth changes and amplitude changes of water, NAA and tCr spectral peaks during the stimulation paradigm. Spectral linewidth-matching procedures were used to correct for the line-narrowing effect induced by BOLD. For an increased understanding of spectroscopic BOLD effects and the optimized way to correct them, a 1 Hz line-narrowing effect was also simulated on mouseproton MR spectrum

^1^H-fMRS data acquired using STEAM acquisitions at 9.4T in rats (n=8) upon optogenetic stimulation of the primary somatosensory cortex were used. Data were analyzed with MATLAB routines and LCModel. Uncorrected and corrected ^1^H-MR spectra of simulated and in-vivo data were quantified and compared. BOLD-corrected difference spectra were also calculated and analyzed.

**Results:** Significant mean increases in water and NAA peak heights (+ 1.1% and +4.5%, respectively) were found accompanied by decreased linewidths (−0.5 Hz and −2.8%) upon optogenetic stimulation. These estimates were used for further definition of an accurate line-broadening factor (lb). Usage of an erroneous lb introduced false-positive errors in metabolite concentration change estimates thereby altering the specificity of findings. Using different water scalings within LCModel, the water and metabolite BOLD contributions were separated.

**Conclusion:** The linewidth-matching procedure using a precise lb factor remains the most performant approach for the accurate quantification of small (±0.3 μmol/g) metabolic changes in ^1^H-fMRS studies. A simple and preliminary compartmentation of BOLD effects was proposed, which will require validation.

## 1. INTRODUCTION

The investigation of neurochemical changes during brain activity is important for the improved interpretation of neurovascular coupling mechanisms in the healthy and diseased brain [1,2]. In this context, proton functional MR spectroscopy (^1^H-fMRS) represents a technique of particular interest when investigating the underpinnings of functional MR imaging signals [1,2] both in the human brain and in animal models. ^1^H-fMRS techniques generate more and more interest allowing for the reproducible assessment of activity-induced changes of important neurotransmitters concentrations such as glutamate (Glu) and g-aminobutyric acid (GABA) as well as energy metabolites such as Glucose (Glc) and Lactate (Lac) [1, 3]. Interestingly, absolute functional changes of these metabolite concentrations demonstrated to be rather small (~0.2μmol/g) in humans [4, 5, 6]. The absolute quantification of these changes was performed following the correction of T_2_*-induced effects [7].

T_2_*-induced effects (or BOLD effects) induce a narrowing of spectral peak linewidths as well as T_2_ changes, as a result of the increased oxygenation within the activated region of interest. These decreases were mainly observed on water, N-Acetyl-Aspartate (NAA) and total Creatine (tCr = PCr+ Cr) singlets and represented only 2% change during visual stimulation of the human visual cortex. They were more easily observable as relative increases in metabolite peak height of around 3% [7]. Although the change in metabolite concentration due to T_2_*-induced effects is low (less than 1 %), if left uncorrected, it may lead to a high degree of false-discovery rate [8]. Errors may become particularly important when absolute changes as low as 0.2 μmol/g are expected. For an accurate estimation of metabolite concentration changes, Mangia et al. [4, 5] proposed to correct for the BOLD effects by line broadening the population-averaged stimulated spectrum to match the linewidth of the corresponding population-averaged REST spectrum. The corrected stimulated and REST spectra were subsequently subtracted resulting in BOLD-free difference spectrum. Positive Glu and Lac peaks were visually identified and subsequently quantified using a simulated difference basis set within LCModel [9]. This procedure demonstrated reproducible outcome within the human primary visual cortex at a high magnetic field owing to the large SNR available [4,5,6,8,10,11,12].

In rodents, ^1^H-fMRS studies remain challenging and have shown lower quantitative reproducibility [13, 14, 15]. The methodology for obtaining accurate estimates of metabolite concentration changes during brain activation remains difficult to replicate in animal models with a voxel of interest more than 500 times smaller than in the human brain [13, 14, 15, 16]. The advent of revolutionary techniques such as optogenetics and chemogenetics that can be coupled to ^1^H-fMRS [16] for the specific stimulation of excitatory cell populations could increase its potential if accurate quantification of metabolic concentration changes can be achieved.

While the difference-spectrum procedure is amplitude-based and pretty straightforward, the line broadening procedure followed by LCModel quantification is ambiguous because areas under the spectral peaks are calculated. On that account, peak line broadening should not affect metabolite quantification. This is, however, not the case. Such errors in metabolite fitting models using linear combination modelling algorithms (such as LCModel) have been scarcely reported in the literature but interest is growing as more accuracy of metabolite concentration estimates and more standardization across processing methods are needed [17, 18]. Besides, these algorithms do not account for the change of the apparent T_2_ value.

In ^1^H-fMRS studies at high magnetic field strength, it is of paramount importance to remove BOLD effects or at least to determine how they affect neurochemical profiles. In human studies at 7T with high SNR levels, corrections were applied and deemed adequate as they compared well with results from difference spectrum procedures [8]. How were these corrections optimised ? How was it decided they were suitable and how can they be appropriately applied to data with poorer SNR ?

In the present paper, BOLD effects and their correction were examined with simulated data and on *in-vivo* proton MR spectra acquired in rats during photo-stimulation of the primary somatosensory forelimb cortex (S1FL). The aim of this preliminary work was to investigate the robustness of the line-matching procedure to correct for BOLD effects and the validity of this corretion within LCModel. In addition, a simple method to better isolate and separate BOLD contributions from metabolite concentrations was proposed.

## 2. METHODS

### 2.1.1 Simulations

A highly resolved ^1^H-MR spectrum acquired at 9.4T with a STEAM sequence and a mouse cryoprobe (Bruker Biospin GmbH, Ettlingen, Germany) in the thalamus of a healthy mouse was used to represent a stimulated ^1^H MR spectrum (STIM). This ^1^H MR spectrum was line-broadened (lb=1 Hz) so that the difference in spectral amplitude of NAA represented a 2% change according to literature values and represented the REST ^1^H MR spectrum (REST). White noise was added to compensate for the smoothing effect. The difference in NAA amplitude was assumed to represent the line narrowing effect induced by the BOLD effects. Metabolite concentrations were quantified using LCModel using two reference standards : (a) the internal water signal from the unsuppressed water scan acquired in the same mouse, which was assumed to represent the BOLD contaminated water signal (for clarity purpose, it was called waterBOLD) and (b) the internal water signal from the unsuppressed water scan acquired in the same mouse and line broadened (to represent a 2% signal decrease compared to waterBOLD), which was assumed to represent the water signal at REST (waterREST).

### 2.1.2 Animals and surgery

All experiments were performed according to the German Tierschutzgesetz and were approved by local authorities (Landesamt für Natur, Umwelt und Verbraucherschutz Nordrhein-Westfalen, Germany). 8 Female Fisher rats (F344) were considered in the present study. Each of them underwent two craniotomies above the primary somatosensory forelimb cortex (S1FL) for the injection of a viral construct [19] and the optic fibre (OF) implantation on the day of MR acquisitions. An OF of 200 μm in diameter was inserted only superficially above S1FL at a depth of 100μm and glued to the skull. Each rat was intubated under isoflurane anaesthesia in oxygen (2-2.5%) and one of the caudal veins was catheterized for infusing pancuronium bromide (1mg/ml BW) during fMRI-fMRS acquisitions. The rat skull was covered with warm agarose gel (1%) to further prevent air-tissue susceptibility artefacts. After fixing the rat head using ear and bite bars, the rat was positioned in a dedicated cradle. Respiration was monitored throughout the entire duration of the experiments as well as end-tidal CO_2_ measurements performed through a capnometer (CapStar-100 CO_2_ Analyzer, CWE Inc., Ardmore, PA, USA) and maintained between 2.8 and 3.5%. The temperature was measured through a rectal probe and maintained at 37±1°C via water tubing linked to a temperature retro-controlled bath.

### 2.2. Magnetic Resonance Imaging and spectroscopic experiments

#### 2.2.1 BOLD-fMRI

All MR experiments were performed at 9.4T (Biospec 94/20, Bruker Biospin GmbH, Ettlingen, Germany) in a small animal MR scanner equipped with 0.7m/T gradients using a 20mm-single loop surface coil for the reception and a 90-mm volume coil for transmission (Rapid Biomedical gmbH, Rimpar, Germany). After the acquisition of pilot and anatomical images, anaesthesia was switched to medetomidine (Domitor, Pfizer, Orion Corporation, Espoo, Finland) (0.04mg/kg (bolus) + 0.05mg/kg/hr (subcutaneous infusion)) and isoflurane was discontinued. The first single-shot gradient-echo echo-planar imaging (TR/TE= 1000/18ms; FOV=28 x 26mm; Matrix= 80 x 80; Bandwidth=200-300 kHz; TH=0.8mm; 16 slices; 600 images) took place at least 1 hour after the start of medetomidine infusion and after whole-brain shimming using MAPSHIM. Laser pulses were delivered successively using in-house developed programs allowing triggering and repetition of a 10sOFF-10sON-10sOFF paradigm (pulse frequency of 9Hz; pulse duration of 10ms). The power calibrated green laser light (552nm) was delivered ensuring a mean power intensity below 22mW/mm^2^ at the tip of the OF to avoid heat effects [20]. BOLD-fMRI images were processed using SPM12 as described earlier^13^.

#### 2.2.2 functional Magnetic Resonance Spectroscopy (fMRS)

Localized functional proton MR spectroscopy was conducted using a STEAM sequence (TR= 4 s ; TE=2.6 ms ; TM= 10ms SW=4960 Hz ; 4096 points). An 8 μl VOI was positioned onto the anatomical T_2_-weighted RARE images coregistered to the GRE-EPI images ensuring that most of the VOI encompassed the BOLD responses while lipid contamination from the skull was minimized. The water signal was suppressed using the VAPOR module [21] and three modules of outer volume saturation (OVS) were interleaved with the water suppression pulses. A 3 x 3x 3 mm^3^ voxel was placed over the rat cortex and used for shimming down to a water linewidth of 15 Hz using first and second-order FASTMAP shimming. A 2.5 min OFF-5min ON-2.5min OFF paradigm was used representing 45 minutes of acquisition (650 FIDs) and repeated 4 times per rat. Unsuppressed water spectra were also acquired using the same sequence and paradigm to provide reference water peaks for eddy current correction and further metabolite quantification. STEAM ^1^H MR spectra were reconstructed and preprocessed using in-house written MATLAB routines. For each rat, raw ^1^H spectra were corrected for frequency drift and FIDs were summed across stimulation and rest periods, respectively, and transferred to a SUN station for LCModel analysis [9] using a basis set provided by Steven Provencher containing a simulated set of macromolecules as well as the following metabolites: Scyllo-Inositol: Scyllo; Alanine: Ala; Aspartate: Asp; Glycero-phosphocholine: GPC; Phosphocholine: PCh; Creatine: Cr; Phosphocreatine: PCr; γ-aminobutyric acid: GABA: Glucose: Glc; Glutamine: Gln; Glutamate: Glu: Glutathione: GSH; myo-inositol: Ins; Lactate: Lac; N-acetyl aspartate: NAA; N-acetylaspartyl-glutamic acid: NAAG; Phosphatidylethanolamine: PE; Taurine: tau. tCr: total creatine (PCr + Cr); Glx: Glu + Gln; Absolute metabolite concentrations were obtained using unsuppressed water signals as an internal reference assuming a brain water content of 80%. The Cramer-Rao Lower bounds (CRLB) were used as a reliability measure of the metabolite concentration estimates. Only metabolites CRLB below 15% were kept for further analysis.

### 2.3 Data analysis

#### 2.3.1 Estimation of T_2_*-induced effects on NAA and tCr spectral peaks

A moving average was used to calculate NAA and tCr signal time courses. Each metabolite concentration-time point was obtained by summing four consecutive blocks of eight FIDs. The procedure was applied to each of the four functional MR spectroscopic acquisitions obtained during the delivery of green laser paradigms. Time courses were then averaged across the animal population. The temporal resolution was 42 s. Unsuppressed water signal time courses were obtained using a 4s temporal resolution for each rat and averaged over the population of rats.

#### 2.3.2 Estimation of metabolite concentration changes

After phasing and correction for B0 shifts, 1200 FIDs were summed per animal and then across 8 rats for the 5-min rest periods and the 5-min stimulation periods resulting in a STIM spectrum and a REST spectrum. STIM and REST spectra were then individually fitted using LCModel. Metabolite concentrations were obtained using either the temporally averaged unsuppressed water peak obtained during resting periods or temporally averaged unsuppressed water peak obtained during stimulation periods. The latter procedure was termed the WATERBOLD approach as described in the theory section.

STIM and REST spectra were also subtracted from each other. BOLD-corrected STIM spectra (STIMC) were obtained using a line-broadening factor lb. lbs were increased in steps of 0.2 Hz and applied to the NAA peak of the averaged stimulated spectrum to match the linewidth of the NAA peak of the averaged REST spectrum. The lb value that best minimized the residual NAA difference peak between STIM and REST was used. STIMC spectra were LCModel fitted.

All procedures were performed using an in-house written Matlab routine. BOLD-corrected difference spectra also obtained (STIMC-REST) and fitted using LCModel after simulation of a basis set containing positive lactate and glutamate changes and negative aspartate and Glucose changes.

### 2.4. Statistics

A Shapiro-Wilk test was performed for all the metabolite concentrations quantified during REST and stimulation conditions demonstrating normality of all the distributions with a p-value above 0.05. Repeated measures One-way ANOVA tests followed by Bonferroni posthoc tests (11 metabolites) to correct for multiple comparisons were used to compare changes in metabolite concentrations. Results were presented as mean ± CRLB. A p-value under 0.05 was considered significant.

Averaged per cent changes in signal time courses between REST and stimulation periods were compared using a paired student t-test. A p-value under 0.05 was considered significant. Results were presented as mean ± standard deviation.

## 3. RESULTS

### 3.1 Simulations of BOLD responses obtained with fMRI and fMRS

#### 3.1.1 Comparison of BOLD responses

For a better understanding of the relationship between T_2_* and spectral peak linewidths during localized brain activation, equations relating to these entities were used.

The full width at half maximum (FWHM) of a spectral peak obeys the following equation:

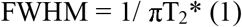

Changes in R_2_*due to focal cerebral activity can be expressed as :

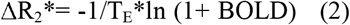

allowing the change in linewidth (Δlw) occasioned by the BOLD effects to be expressed as:

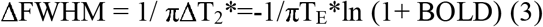

These equations were used to model and compare « BOLD-fMRI» and «BOLD-fMRS» responses on water.

As the magnetic field strength increases, the T_2_* relaxation time decreases significantly [22]. Upon BOLD activation of the human motor cortex, changes in relaxation rate ΔR_2_* increased linearly as a function of field strength [23]. A higher spectral peak linewidth change due to activation should therefore be expected at higher magnetic field strength for a same TE. The assessment of T_2_*-induced effects due to hyperoxygenation of blood should be easier at high magnetic field strength (> 7T).

Water linewidth changes (Δlw) for TE = 3, 18, 26 ms, chosen from previous ^1^H-fMRS studies, were calculated with Eq. (3) (**Fig. 1**). Δlw changes evolved linearly with BOLD changes. At TE= 3 ms, changes were elevated compared to those obtained at TE= 18 ms (9.4T, [16] or TE= 26 ms (7T, [4, 5]).

**Figure 1.**
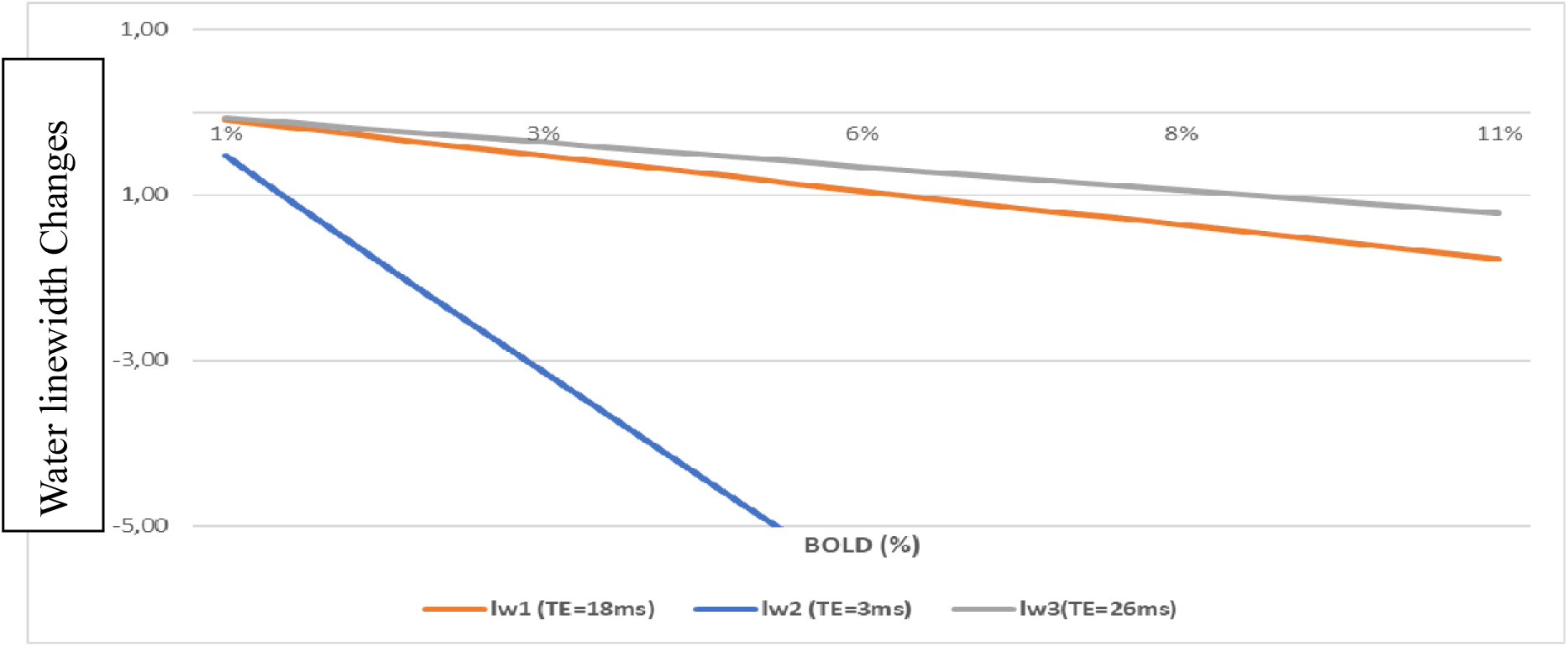
Water linewidth changes versus BOLD-fMRI for different TEs using equations (1) and (3) However for TE= 3 ms, the BOLD effects measured with fMRI is negligible. For ΔT_2_*= 1.5 s (arbitrarily chosen), BOLD = 0.47% for TE= 3 ms when BOLD = 2.9 % for TE = 18 ms. This change corresponds to a linewidth change of - 0.5 Hz and BOLD = 0.5% for TE = 3 ms and BOLD = 3 % for TE = 18 ms. These values correspond well to linewidth changes quantified *in vivo* on NAA and Cr spectral lines [10,24].

For BOLD = 2% Δlw =-2.1 Hz (TE=3 ms) ; Δlw =-0.35 Hz (TE=18 ms) ; Δlw =-0.24 Hz (TE=26 ms) ;

#### 3.1.2 Quantification of metabolite concentrations affected by simulated BOLD effects

To illustrate errors induced by the BOLD effects and an erroneous correction of this effect on metabolite quantification, a 2% BOLD effect was simulated that induced a 1 Hz decrease of the NAA spectral peak linewidth (Fig 2). Metabolite concentrations were quantified using LCModel [9]. In this example, no metabolic concentration changes occurred. The ground truth metabolic concentrations were obtained by LCModel adjustment of the non-stimulated spectrum (black spectrum).

**Figure 2:**
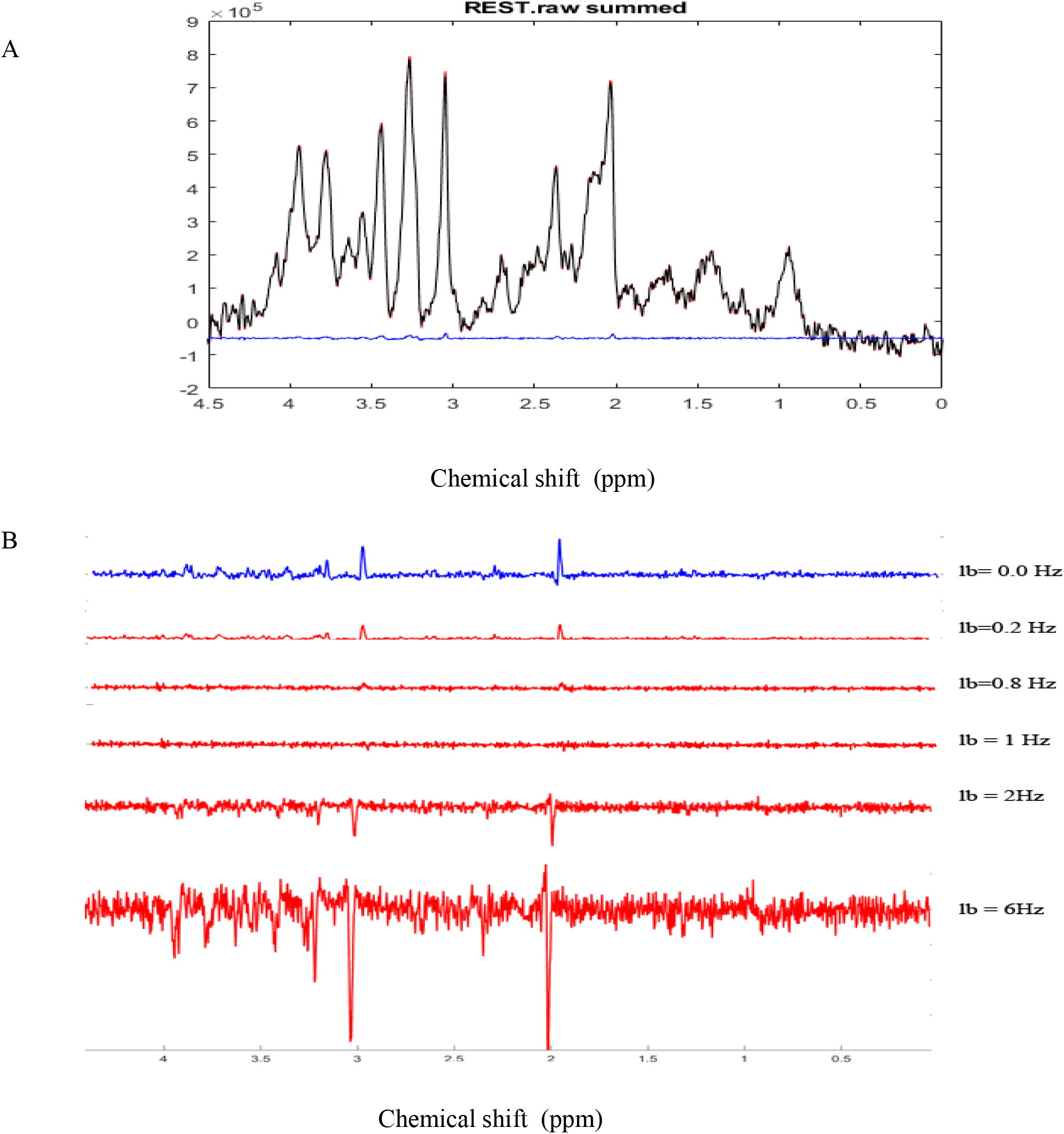
Simulation of BOLD effects : A. A highly spectrally resolved ^1^H-MR spectrum acquired at 9.4T in the mouse thalamus with a cryoprobe (red) served as a stimulated spectrum (STIM) and was exponentially line broadened by 1 Hz to represent a 2% amplitude decrease of the NAA peak in the resting spectrum (REST, black). The 2% difference represents the simulated BOLD effect as a result of T_2_*-induced effects. BOLD effects can also be seen as residual positive peaks in the difference spectrum (blue; STIM-REST) B. C. The REST spectrum was subtracted from the STIM spectrum demonstrating residual positive peaks due to the simulated BOLD effect. As the line-broadening factor lb applied to the STIM spectrum increased from 0.2Hz to 1Hz, BOLD residuals gradually disappeared while negative residuals grew up for lb values above 1 Hz

The correction for BOLD effects consists of the application of a line broadening factor, which must be at least equal to the decrease of the spectral linewidth induced by focal cerebral activity so that stimulated and non-stimulated spectral linewidths are equivalent. This correction can be applied to the water peak [7] but most ^1^H-fMRS studies report it on N-acetyl-aspartate (NAA) and total Creatine (PCr + Cr) peaks [4, 5, 6, 8, 13]. To observe changes induced by the BOLD effects, MR spectra obtained during stimulation and REST are subtracted. Residuals can be visually observed and correspond to the BOLD effects (**Fig. 2**).

The corrected neurochemical profiles of the spectra depicted in Fig. 2 are displayed in Fig 3. REST represents the neurochemical profile for «true» metabolite concentrations while STIMC represents the corrected neurochemical profile for which a 1 Hz line broadening correction was applied. This correction represents the optimum correction for BOLD effects since metabolite concentration differences were minimized. The line broadening correction is the most effective when it is the closest to the linewidth reduction induced by the BOLD effects. The error relative to REST remained below 1% for lb= 1 Hz for all the metabolites. The error rose to 4 % for lb= 6 Hz for PCr and Ins and was of the same order of magnitude as the error induced by the non-corrected BOLD effects (lb=0 Hz).

**Figure 3:**
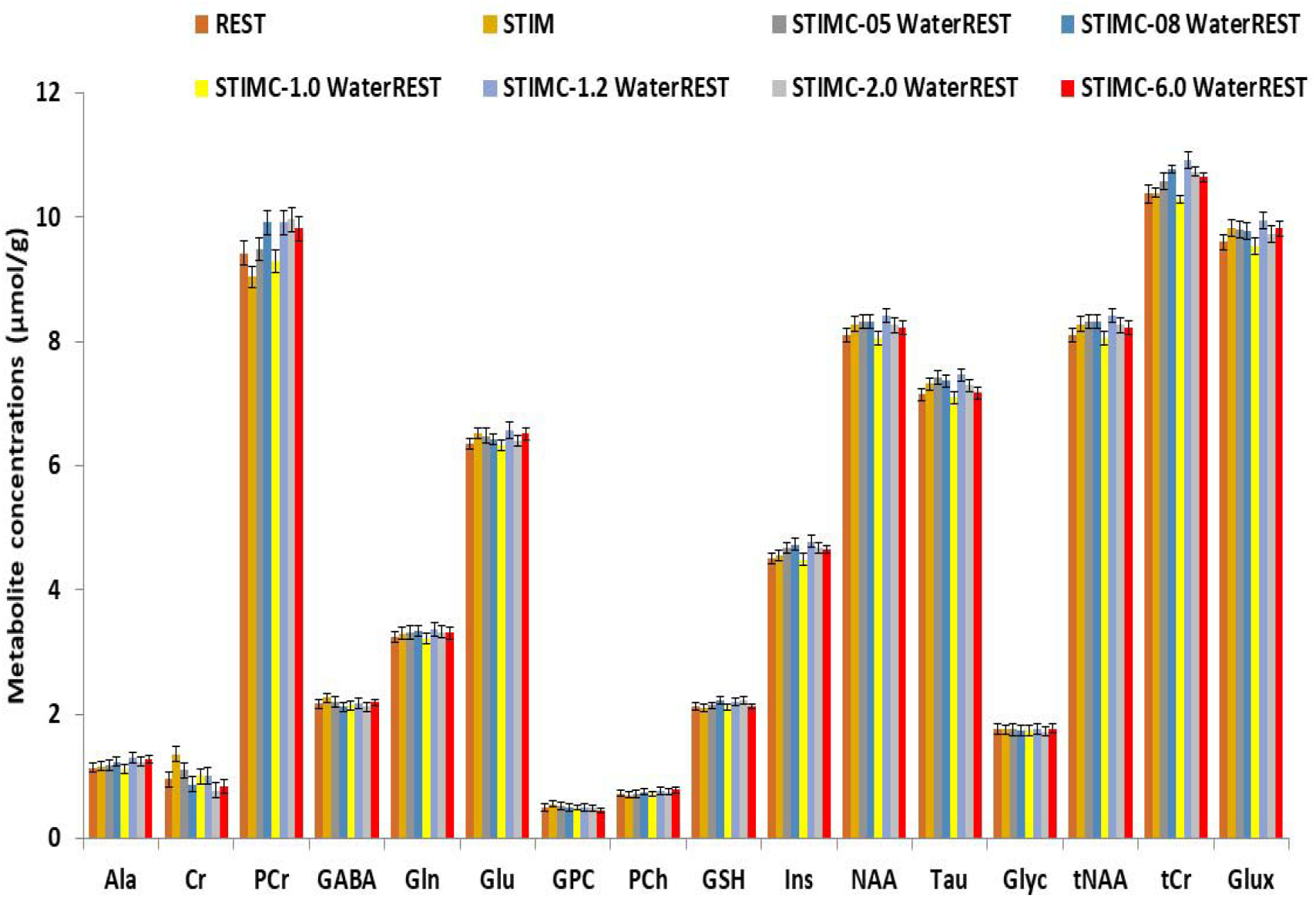
Comparison of neurochemical profiles. REST represents the ground truth for metabolic concentrations. Line broadening (lb) factors of 0.5 Hz, 0.8 Hz, 1 Hz, 1.2 Hz, 2 Hz and 6 Hz were applied to the simulated STIM spectrum. The correction that best approached the ground truth was for lb= 1 Hz.

In Fig 4, STIM represents the uncorrected BOLD effect neurochemical profile. The quantification was performed with waterREST, the unsuppressed water spectrum acquired before the REST spectrum. BOLD effects increased Glu concentrations by 2.6 % and NAA concentrations by 2.2 %. «waterBOLD» represents the neurochemical profile acquired during stimulation but quantified using the simulated unsuppressed water peak with BOLD effects. Table 1 compares the Glu, NAA and tCr concentrations quantified with LCModel for the different cases. When using waterBOLD for quantification, the BOLD effects increased Glu concentrations by 1.8 % and NAA concentrations by 1.4 %. The error induced by BOLD effects, BOLD_water_, on the water was calculated as the metabolite concentration difference between STIM and waterBOLD and was approximately 0.8 % for all metabolites. The error induced by BOLD effects, BOLD_metab_, on the water was calculated as the metabolite concentration difference between STIM and STIMC. Interestingly, BOLD effect-induced errors were cumulative: BOLD_STIM_ = BOLD_metab_ + BOLD_water_. Therefore, it is postulated that estimates of metabolite concentrations with waterBOLD are affected by BOLD on metabolites only.

**Figure 4:**
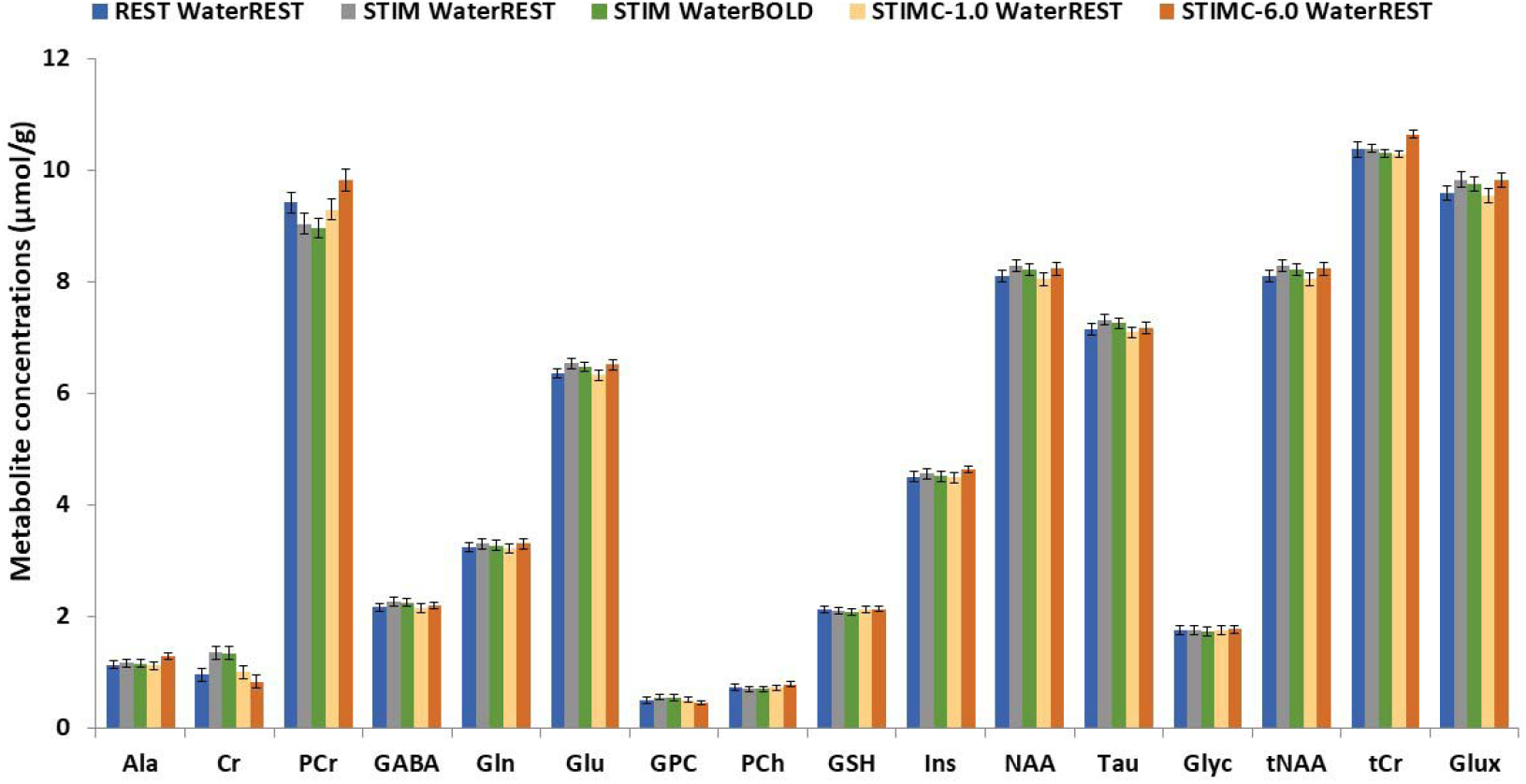
Comparison between the REST, STIM, STIMC and waterBOLD neurochemical profiles. STIMC is the neurochemical profile of the line-broadened STIM spectrum with lb = 1 Hz. waterBOLD is the neurochemical profile of the STIM spectrum scaled with the stimulated water peak.

**Table 1.** LCModel quantification of Glu, NAA and tCr (± CRLB) for simulated REST and STIM spectra and STIM scaled with waterBOLD and the line-broadened STIM spectrum (STIMC lb = 1 Hz)

|  | REST | STIM | waterBOLD | STIMC |
| --- | --- | --- | --- | --- |
| Glu ($\mu\text{mol/g}$) | 6.36±0.1 | 6.5±0.1 | 6.48±0.1 | 6.32±0.1 |
| NAA ($\mu\text{mol/g}$) | 8.11±0.1 | 8.3±0.1 | 8.22±0.1 | 8.05±0.1 |
| tCr($\mu\text{mol/g}$) | 10.38±0.1 | 10.4±0.1 | 10.31±0.1 | 10.29±0.1 |

### 3.2 *in-vivo* characterization of BOLD effects

In the rat S1FL, optogenetic stimulations resulted in an average BOLD response measured with fMRI of 2.4± 1.3 % (Fig.5 A). A typical T-value BOLD map overlaid onto an anatomical image is shown (**Fig. 5A**) as well as a typical BOLD timecourse following a 10-minute (10s ON-20s OFF) stimulation paradigm (**Fig. 5B**). Representative raw ^1^H MR stimulated (**Fig. 5C**) and REST (**Fig. 5D**) spectra acquired during the same paradigm of stimulation were acquired. The mean relative linewidth changes in NAA (Δlw_NAA_) were −2.8 ± 4 % (n=8) and the mean relative tCr linewidth changes (Δlw_tCr_) were −1.4 ± 4% (n=8).

**Figure 5:**
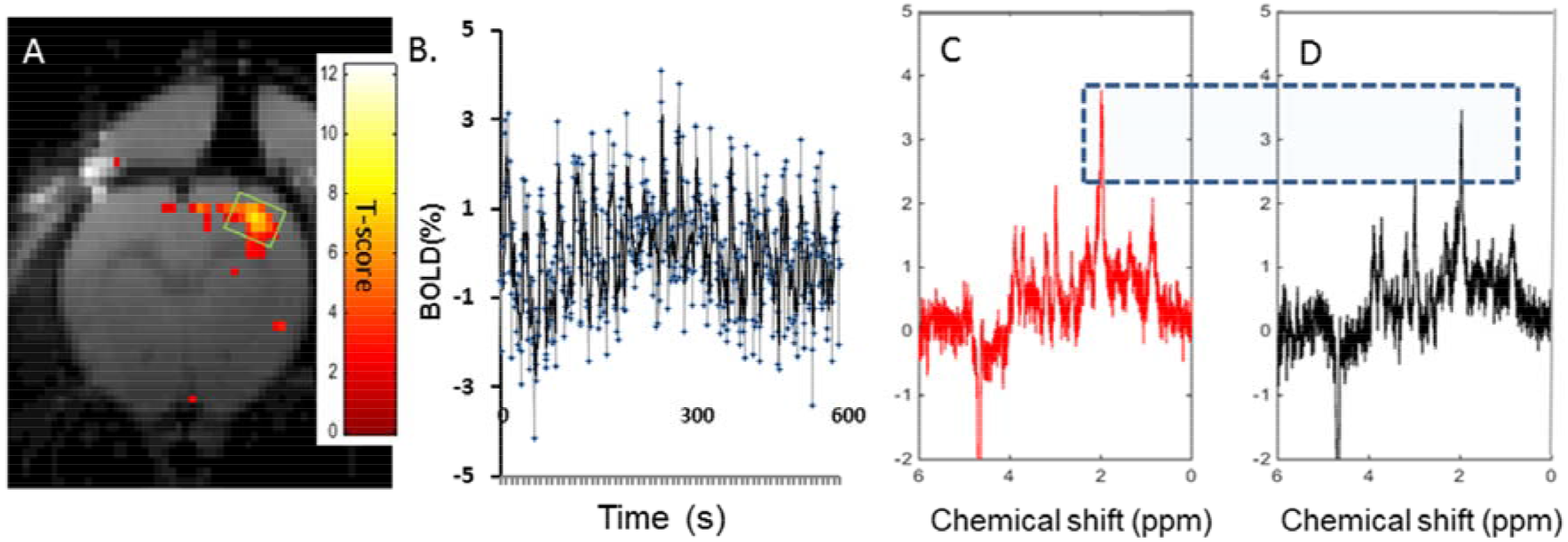
Characterization of BOLD effects on and tCr signals : A. Typical T-value BOLD map overlaid onto an anatomical image of the rat brain demonstrating activation in S1FL in a rat upon optogenetic stimulation. B. Typical BOLD timecourse during a 10s-ON-20s OFF paradigm of stimulation lasting 10 minutes. C and D. Single rat ^1^H-MR spectra acquired in a 8 μl voxel of interest covering the activated NAA area (as shown in Fig. 5A) during optogenetic stimulations (325 FIDs)(C, red) and resting periods (D, black). Boths spectra were reconstructed from FIDs acquired during the same paradigm of stimulation (5min-ON-5-min OFF, 45 minutes). The blue dotted rectangle serves to show the slightly increased amplitude of NAA in the stimulated spectrum compared to the resting spectrum.

For an improved characterization of the BOLD effects on ^1^H MR spectra, water, NAA and tCr signals were measured over time during optogenetic stimulation paradigms (5 min ON-5min OFF). NAA peak height followed the optogenetic stimulation paradigm (**Stimulation vs. REST vs. Stimulation**: 8.4 ± 6% vs. 3.2 ± 4.3% (P >0.05) vs. 4.5± 4.4%; P<0.05; Paired student t-test; **Fig. 6A**). The relative change of tCr peak height did not follow the paradigm of stimulation (**Fig. 6B)**.

**Figure 6:**
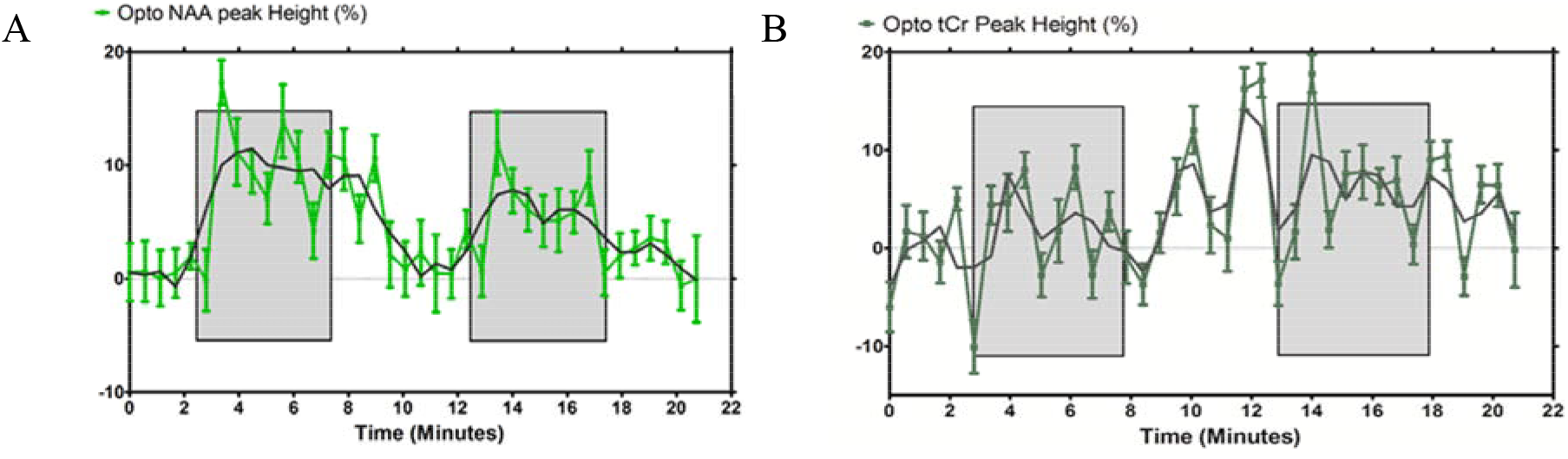
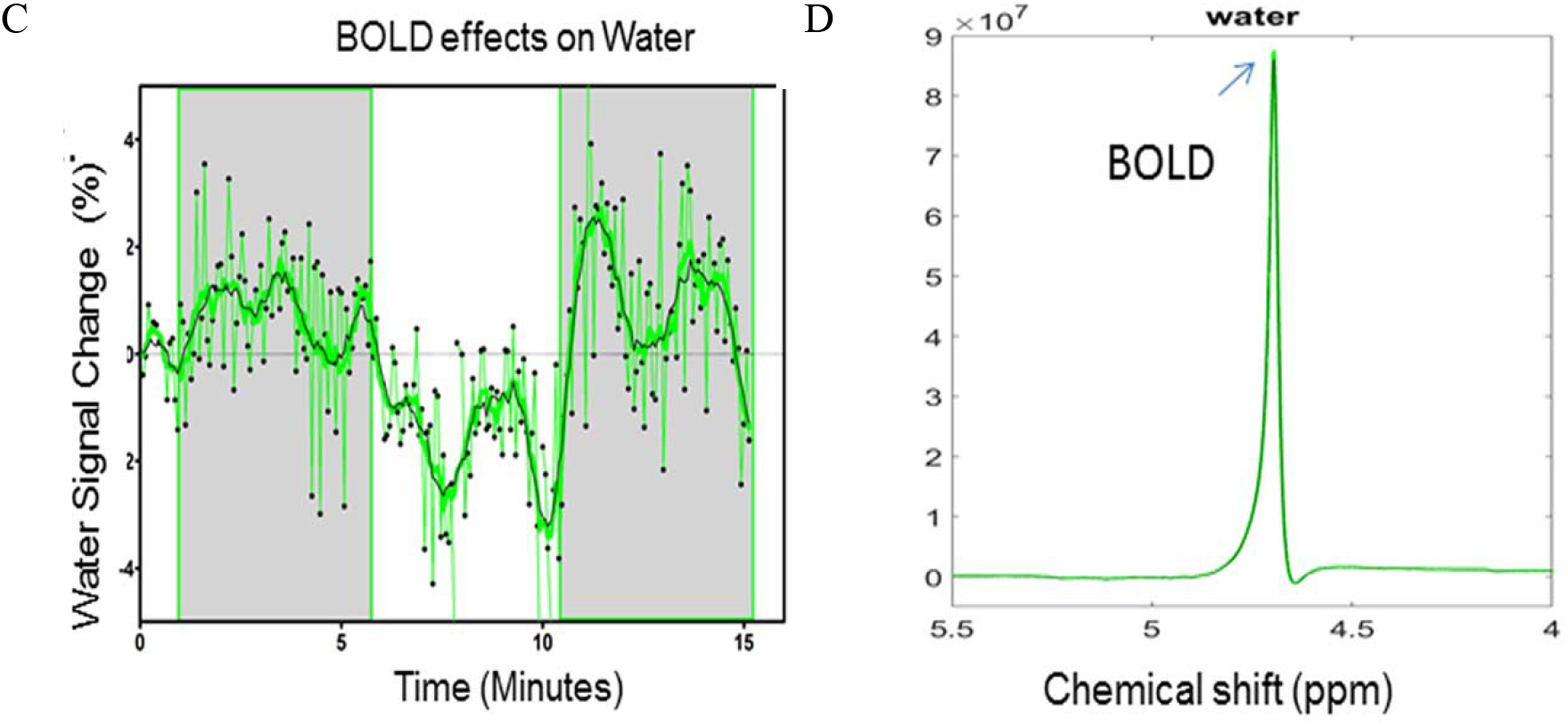
**A**. Quantification of NAA and **B**. tCr relative peak height changes (% ± s.d) for optogenetic stimulations Shaded areas indicate the periods of stimulation. Statistical comparisons are described in the text. **C**. Quantification of water peak height changes (%) for optogenetic stimulations during the paradigm. Shaded areas indicate periods of stimulation. **D**. The unsuppressed water peaks were temporally averaged for resting periods (black) and stimulated periods (green) demonstrating a difference in amplitude (arrow) that can be attributed to BOLD effects. Averaged water peaks were both used for further quantification of metabolite concentrations.

Unsuppressed water signals were also acquired during the paradigm of stimulation as a function of time (n=5) **(Fig. 6C)**.

Green laser stimulation induced significant increases in water peak heights over 5 minutes of S1FL activation relative to rest periods (REST vs. Stimulation vs. REST vs. Stimulation: 0.1± 0.7% vs. 0.8±1.2 % (P=0.013) vs −1.6±1.4 % (P<0.001) vs. 1.1± 1.5 (P<0.001), Student t-test; Fig.6C; mean ± standard deviation). Stimulated and resting water peaks were averaged into single water peaks shown overlaid in Fig. 6D. The mean relative linewidth changes in water (Δlw_water_) were −0.5 ± 0.8%. Mean peak height and linewidth changes between REST and stimulation are summarized in Table 2.

**Table 2.** *In-vivo* relative percent change of the height and linewidth of Water, NAA and tCr peaks (optogenetic versus REST)

| % Height Change |  |  | % Linewidth Change |  |  |
| --- | --- | --- | --- | --- | --- |
| Water | NAA | tCr | Water | NAA | tCr |
| 1 ± 1.0 | 3.3 ± 7 | NA | -0.5±0.8 | -2.8 ± 4 | -1.4 ± 4 |

### 3.3 *In-vivo* quantification of metabolite concentration changes affected by BOLD effects

Population-averaged ^1^H MR spectra (n=8) acquired during optogenetic stimulation and REST are depicted in **Fig.7a** and **7b** respectively. The REST spectrum was subtracted from the stimulated spectrum resulting in a difference spectrum (**Fig. 7c**). The NAA peaks of the stimulated and REST spectra were line-matched using a line broadening factor of 0.5 Hz. The lb factor was gradually increased in steps of 0.2 Hz up to 1 Hz by minimizing the residual NAA difference peak (**Inset)**. Each ^1^H MR spectrum was individually fitted using LCModel. Their subtraction resulted in a BOLD-corrected difference spectrum (**Fig.7d**). An identical line-matching procedure was performed using lb= 6 Hz corresponding to a 6 % change in NAA peak height during optogenetic stimulation. The 6 Hz line broadened spectrum was also LCModel fitted. Metabolite concentrations quantified using LCModel and their changes are summarized in Table 3.

**Figure 7:**
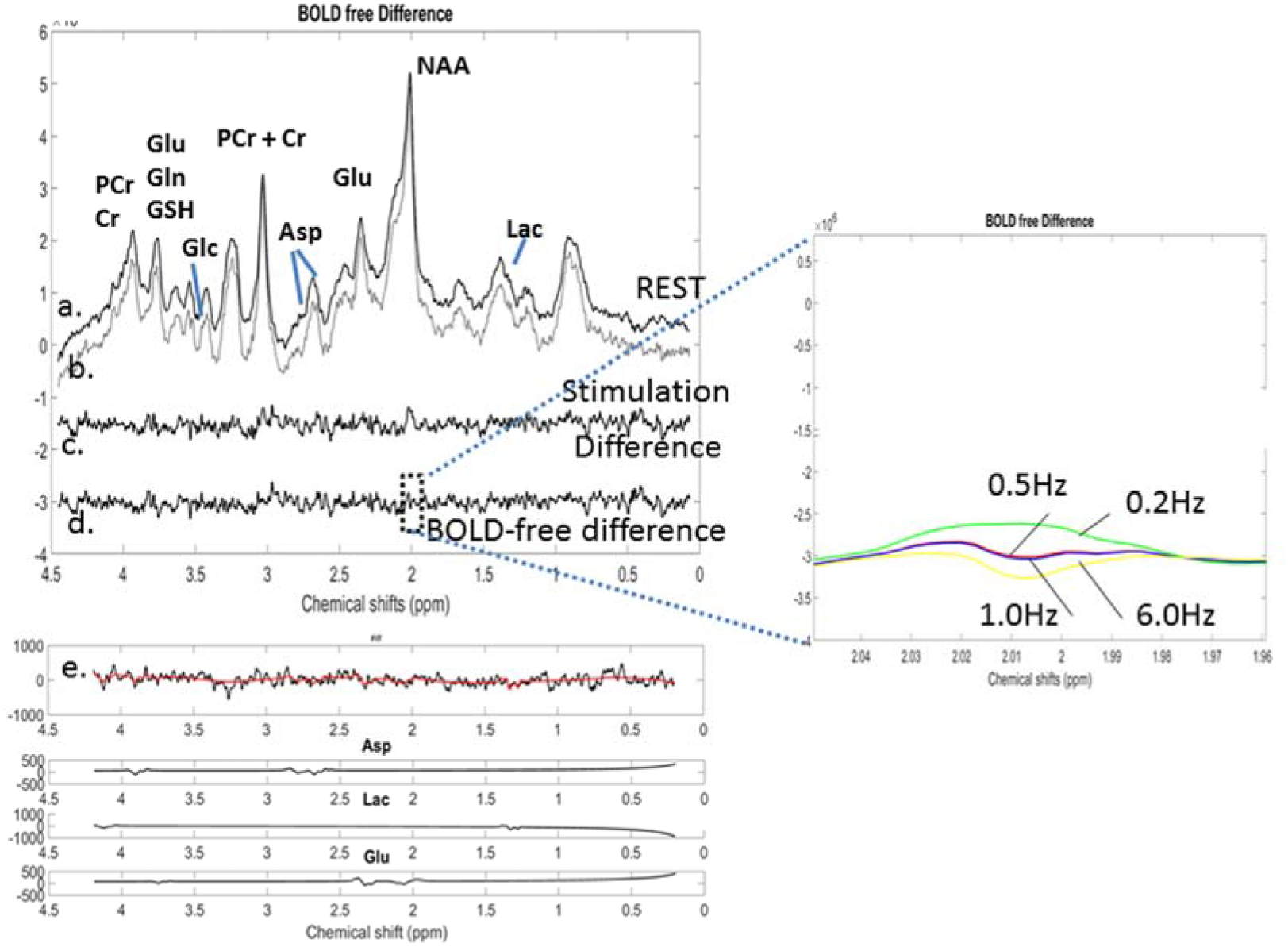
o-fMRS: Averaged (n=8) and labeled proton MR spectra for a. resting conditions and for b. Photo-stimulation with green light. c: The REST spectrum was subtracted from the stimulated spectrum prior to any correction for BOLD effects d. The BOLD free difference spectrum was obtained following application of a 0.5 Hz line broadening to minimize BOLD effects. e. LCModel fitting of the BOLD free difference spectrum allowed identification of the polarity of metabolic changes and retrieved Asp, Lac and Glu components included in the simulated basis set of the BOLD-free difference spectrum. Inset: NAA residuals for lb=0.2 Hz, 0.5 Hz, 1 Hz and 6 Hz resulting from the subtraction of the REST spectrum from line-broadened stimulated spectrum. The NAA residual was best minimized for lb= 0.5 Hz

**Table 3:** Quantification of *in-vivo* metabolite concentrations: Baseline, Stimulated and BOLD corrected metabolite concentrations were obtained through adjustments of MR spectra with LCModel. Stimulated MR spectra were quantified using waterREST and WaterBOLD respectively. The optimized line broadening factor (see Fig.7) was obtained for lb= 0.5 Hz. An erroneous lb = 6 Hz was also used. A group analysis was performed and one way ANOVAs were performed to evaluate the significance of metabolite concentration changes. Bonferroni post-hoc tests were used for multiple comparisons. * p< 0.05; ** p< 0.01; *** p< 0.001

|  | Baseline concentration |  | STIMULATED |  |  |  |  |  |  |  | BOLD CORRECTED |  |  |  |  |  |  |  |
| --- | --- | --- | --- | --- | --- | --- | --- | --- | --- | --- | --- | --- | --- | --- | --- | --- | --- | --- |
|  | REST |  | No linewidth correction |  |  |  | STIM/WaterBOLD |  |  |  | Linewidth matching lb =0.5 Hz |  |  |  | Linewidth matching lb =6 Hz |  |  |  |
|  |  |  | Concentration difference |  |  |  | Concentration difference |  |  |  | Concentration difference |  |  |  | Concentration difference |  |  |  |
|  |  | CRLB |  | CRLB | Mean (μmol/g) | Mean (%) |  | CRLB | Mean (μmol/g) | Mean (%) |  | CRLB | Mean (μmol/g) | Mean (%) |  | CRLB | Mean (μmol/g) | Mean (%) |
| Asp | 4,31 | 10% | 3,13 | 7% | -1,18 | -27,3*** | 3,15 | 8% | -1,16 | -26,8*** | 4,07 | 8% | -0,24 | -5,5** | 4,06 | 6% | -0,26 | -5,9** |
| Gln | 3,79 | 3% | 3,58 | 3% | -0,21 | -5,6* | 3,53 | 3% | -0,26 | -6,9** | 3,58 | 4% | -0,21 | -5,5* | 3,37 | 3% | -0,41 | -10,9*** |
| Glu | 6,98 | 2% | 7,54 | 2% | 0,56 | 8,05*** | 7,37 | 2% | 0,40 | 5,66*** | 7,52 | 2% | 0,54 | 7,7*** | 8,57 | 2% | 1,59 | 22,8*** |
| GSH | 1,44 | 5% | 1,20 | 6% | -0,23 | -16,3** | 1,17 | 7% | -0,27 | -18,6** | 1,51 | 5% | 0,07 | 5,1 | 1,84 | 4% | 0,40 | 28,0*** |
| Ins | 2,91 | 5% | 3,64 | 3% | 0,73 | 25,1*** | 3,82 | 4% | 0,91 | 31,2*** | 3,54 | 6% | 0,63 | 21,6*** | 3,58 | 3% | 0,66 | 22,8*** |
| NAA | 8,70 | 2% | 9,00 | 2% | 0,30 | 3,5*** | 8,71 | 2% | 0,01 | 0,14 | 8,47 | 2% | -0,23 | -2,7** | 8,85 | 1% | 0,15 | 1,76 |
| Tau | 4,44 | 3% | 4,33 | 2% | -0,11 | -2,5 | 4,26 | 3% | -0,18 | -3,94 | 3,97 | 3% | -0,47 | -10,5*** | 4,14 | 2% | -0,29 | -6,62*** |
| Glyc | 2,05 | 5% | 1,64 | 6% | -0,41 | -19,9*** | 1,35 | 8% | -0,70 | -34,3*** | 1,60 | 11% | -0,45 | -21,8*** | 1,44 | 5% | -0,61 | -29,84*** |
| tNAA | 9,44 | 2% | 10,57 | 1% | 1,12 | 11,9*** | 10,67 | 2% | 1,22 | 12,95* | 10,47 | 2% | 1,02 | 10,8*** | 10,85 | 1% | 1,41 | 14,91 |
| tCr | 8,00 | 2% | 7,64 | 1% | -0,36 | -4,5*** | 7,52 | 2% | -0,48 | -6,04*** | 8,18 | 2% | 0,18 | 2,3 | 7,73 | 1% | -0,27 | -3,40** |
| Glx | 10,77 | 2% | 11,12 | 2% | 0,35 | 3,3*** | 10,90 | 2% | 0,14 | 1,25 | 11,10 | 2% | 0,33 | 3,1*** | 10,94 | 2% | 0,18 | 1,64 |

Linewidth-matching techniques resulted in metabolite concentrations dropping by −0.06 ± μmol/g on average for lb=0.5 Hz and −0.18 ± μmol/g for lb = 6 Hz compared to uncorrected concentrations. The number of metabolite concentrations changing significantly due to stimulation differed with each analysis: waterBOLD vs STIMC (lb=0.5 Hz): 9 metabolites out of 11; p<0.01; waterBOLD vs STIMC (lb=6 Hz):4 metabolites out of 11; p<0.01; STIMC (lb=0.5 Hz) vs STIMC (lb=6 Hz): 6 metabolites out of 11; p<0.01).

Relative to STIM, water scaling with waterBOLD induced modest metabolic concentration changes (below ±0.3 μmol/g) whereas the line-matching-techniques induced important concentration changes for Asp and Glu concentrations (~ −1.0 μmol/g). The changes in Glu concentrations were in the range + 0.4 to + 1.59 μmol/g for all BOLD corrections and remained highly significant (Table 3). This was also the case for Asp concentrations ranging from −1.16 to −0.24 μmol/g.

In an identical manner to the previous simulated data, the total BOLD effects measured *in-vivo* can be calculated as the sum of the BOLD effects on water and the BOLD effects on metabolites. As for the simulated data, BOLD effects on water were calculated as the difference between metabolite concentrations quantified on the stimulated MR spectra with waterREST and waterBOLD respectively. Assuming that the line-broadening correction leads to BOLD-free metabolite concentrations, the BOLD effects on metabolites were calculated as the difference between stimulated metabolite concentrations scaled with waterBOLD and these BOLD-free metabolite concentrations. Thus, for NAA assumed as an intracellular metabolite and a neuronal marker:

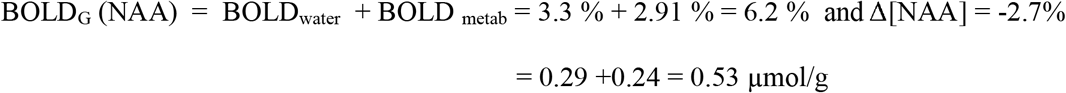

For Glu, the same applies:

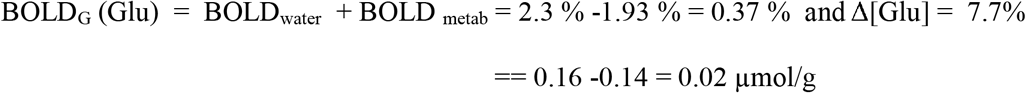

Glu also represents a neuronal marker but is also present in the extracellular space during synaptic release [24]. Exchanges between intracellular and extracellular spaces may explain the negative effect on metabolites and a lesser effect of global BOLD (BOLD_G_). In the same manner, BOLD effects on a glial marker such as Ins, can be separated. The negative contribution of BOLD effects on water may be attributed to changes in water dynamics within astrocytes [25].

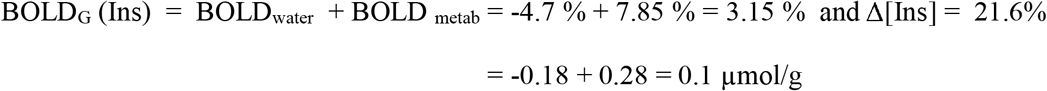

BOLD effects can thus substantially contaminate metabolite concentrations.

### 3.3 Characterization and quantification with BOLD-free difference spectrum

BOLD-free difference spectra were fitted using LCModel and a simulated basis set including Asp, Glu, Glc and Lac **(Fig. 7e)**. The obtained concentration changes for Asp and Glu presented in Table 4 were in agreement with the group analysis performed between averaged STIM and REST spectra (Table 3). A negative Lac peak was also determined. Upon visual inspection of the overlaid optogenetic and REST spectra (**Fig. 8**), positive peaks were observed for Glu (at 2.35 ppm) as well as negative peaks at 3.75 ppm. A negative Asp peak was also identified (2.77ppm). Moreover, positive GSH peaks were also observed (at 2.92/2.97 and 3.78 ppm). At 1.32 ppm, the stimulated Lac peak was under the non-stimulated one resulting in a negative Lac peak in the BOLD-free difference spectrum. In addition, at 3.5ppm, the peak corresponding to Glc disappeared upon optogenetic stimulation, resulting in a negative peak. The quantitative outcome of the LCModel fit to the BOLD-free difference spectrum is reported in Table 4 although noise levels remained important impeding an appropriate and precise quantification of changes using LCModel.

**Figure 8:**
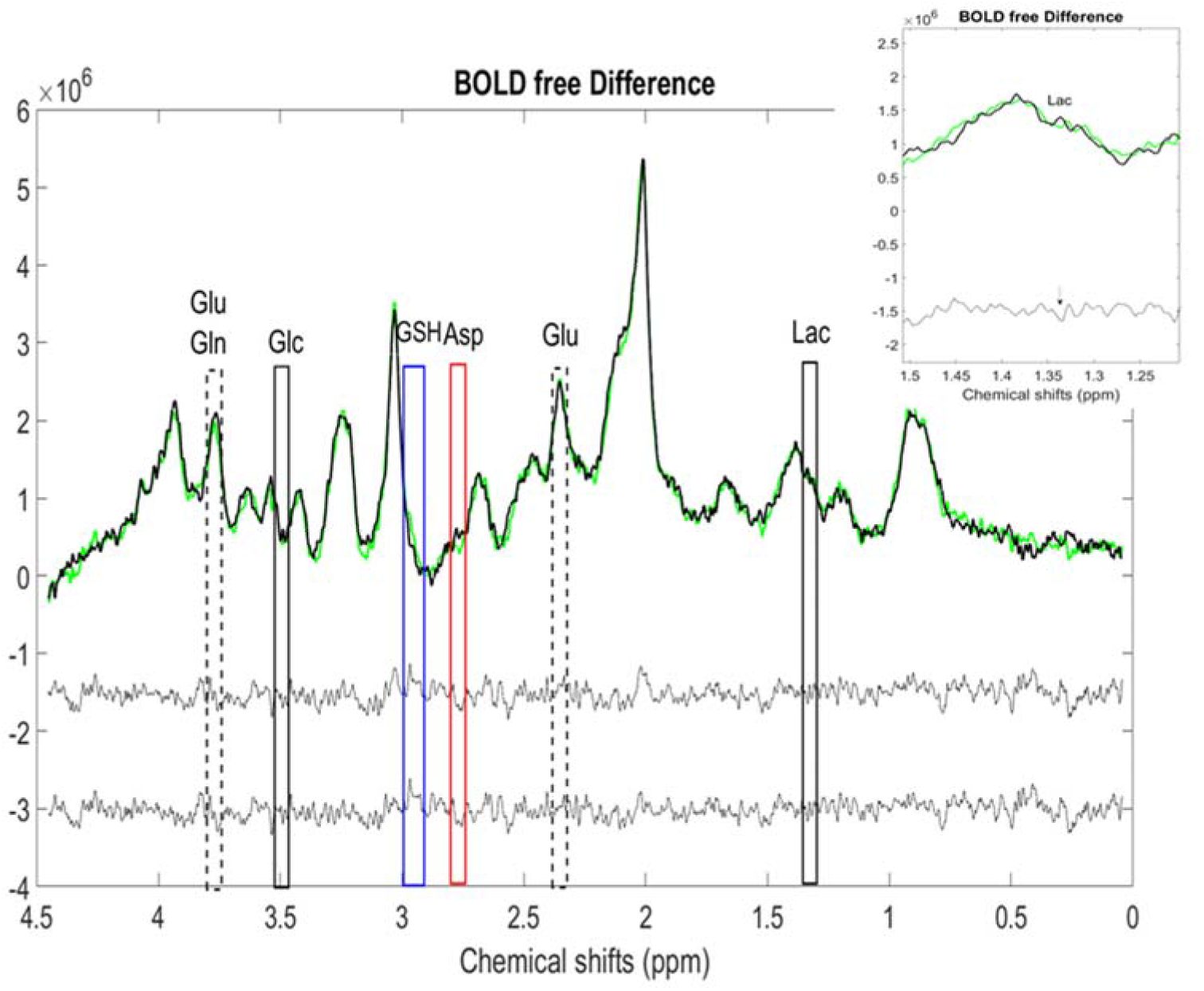
Qualitative observation of amplitude spectral changes: Optogenetically stimulated and REST spectra (n=8) were overlaid allowing enhanced observation of metabolite changes. Overlaid peaks ascribed to lactate (Lac (1.32ppm); plain black rectangle), glucose (Glc (3.5 ppm)), glutamate (Glu (2.12, 2.35 and 3.75 ppm; dotted rectangles)), aspartate (Asp (2.77 ppm); red rectangle) as well as glutathione (GSH (2.92, 2.97 and 3.78 ppm); blue rectangle) were identified. Inset: The arrow indicates a peak of Lac at 1.32 ppm at REST but not during green laser stimulation: amplitude differences lead to a negative peak in the BOLD-free difference spectrum (arrow).

**Table 4:** Quantification of metabolite changes with LCModel

| **Metabolite** | **Concentration Changes ± CRLB ($\mu\text{mol/g}$)** |
| --- | --- |
| **Asp** | 0.3±0.03 |
| **Glu** | 0.46±0.05 |
| **Lac** | 0,8±0.1 |

## 4. DISCUSSION

The present study examined the impact of BOLD effects and their correction on the quantification of metabolites using simulations and *in-vivo* proton MR spectra acquired in the rat primary somatosensory cortex during optogenetic stimulation and rest periods. In rodents, the estimation of reliable metabolite concentration changes due to a functional challenge remains difficult and requires large animal populations [26] and higher magnetic fields or cryoprobes [26, 27] to enhance the SNR.

### 4.1 The line-matching procedure removes BOLD effects

Removal of BOLD effects in ^1^H-fMRS allows for the estimation of metabolite concentration changes solely attributed to neurochemical changes as a consequence of brain activity [1,5,6,8,10,11,12,13,14,28]. BOLD effects generate small changes in metabolite concentrations that are not necessarily of identical magnitude for all metabolites [8, 29]. When not corrected, metabolic concentration changes may appear erroneously significant [12]. The appropriate line broadening correction for BOLD effects is based on the value that best minimizes the residuals of tCr and NAA peaks resulting from the subtraction of the population-averaged stimulated and REST proton MR spectra. This procedure strongly relies on an optimized phasing of both the stimulated and REST spectra to limit potential frequency drifts during the subtraction. Low SNR levels in small VOIs of the rat cortex further complicate the subtraction methodology. A more direct quantification of BOLD-free metabolite concentration changes with LCmodel could simplify ^1^H-fMRS data analysis and interpretation.

Recent studies reported that LCModel analysis of short-TE data was highly sensitive to noise and spectral linewidth variations [30]. Notably, line broadening of original data analyzed with LCModel induced substantial metabolite quantification changes. Therefore, the reliability of metabolite quantification following linewidth matching for BOLD correction can be questioned. In the present study, the simulation of BOLD effects demonstrated that the line-matching procedure is an effective method to remove them with a quantification error of less than 1 % (± 0.05 μmol/g). However, if the line broadening factor is different from the line-narrowing factor induced by BOLD effects, up to a 5 % (± 0.3μmol/g) error in quantification may be obtained, which is substantial when absolute metabolic changes as low as 0.2 μmol/g are expected [4, 5, 6].

### 4.2 Water Scaling

For the absolute quantification of metabolites with LCModel, the unsuppressed water peak acquired in the same VOI as the ^1^H-MR spectrum serves for scaling and for eddy -current correction. During stimulation, the water peak undergoes a line-narrowing effect due to the BOLD effects. As LCMOdel [9] considers resonance areas, the water line-narrowing due to T_2_* -induced effects should not affect metabolite quantification. The simulations used in the present study showed that this was not the case. Using waterBOLD for scaling, the error induced on metabolite concentrations was lower than with waterREST. The quantification of metabolite concentrations on a proton MR spectrum with BOLD simulated contamination suggested that compensatory effects take place when using stimulated water (waterBOLD). This finding led us to postulate that metabolite concentrations quantified using waterBOLD are affected by BOLD effects on metabolites only whereas metabolite concentrations quantified using waterREST contain the cumulative BOLD effects on metabolites and water.

In single-voxel MR spectroscopy, lineshapes can be distorted by B0 field inhomogeneities and eddy currents. Metabolite fitting models using water reference lines to compensate for B0 field inhomogeneity have also been used in combination with LCModel [9] for quantification purposes. Notably, increased accuracy and stability of spectral quantification were found using the reference lineshapes from the water signal [17]. In the present study, the use of waterBOLD for scaling, partially corrected for the change in spectral linewidth occasioned by BOLD effects.

### 4.3 Compartmentation of BOLD effects

BOLD responses measured by fMRI or fMRS on water represent the T_2_*-induced effects encompassing both intravascular and extravascular compartments. Since BOLD fMRI provides an indirect measure of neuronal activity and relies on neurovascular coupling mechanisms, diffusion fMRI (dfMRI) [31] has been proposed as a possible method to disentangle the vascular and extravascular components of BOLD. dfMRI relies on microstructural changes driven by neural activity such as cell swelling to induce changes in the diffusivity of water molecules.

Another technique independent of neurovascular coupling and taking advantage of the specific distribution of metabolites within different tissue compartments is MRS. Neuronal compartments and other extravascular and intravascular compartments may be differentially affected by T_2_*-induced effects. For example, T_2_*-induced effects measured with ^1^H-fMRS on the neuronal markers NAA and Glu and on the glial marker Ins may represent the effects on the neuronal compartment and the glial compartment [7]. In the human visual cortex at 4T, the changes in linewidth and amplitude were similar for water and metabolites [7],which was not the case in the present study at 9.4T. Thus, the linewidth and amplitude changes of the spectral lines assessed with ^1^H-fMRS could represent specific markers of the BOLD effects in extravascular compartments.

Older studies revealed that following injections of the paramagnetic agent (Dy-TTHA)^3-^and a vasodilator, the water peak in the rat brain divided into 3 peaks representing intracellular water, extravascular-extracellular water and intravascular water. The distribution of Na^+^ ions in the 3 different compartments was also shown [32]. When blocking the sodium pump with ouabain, a slight increase of the intracellular water peak signal was noticed corresponding to an increase of the intracellular volume. The intravascular and extravascular-extracellular volumes became almost negligible.

By extrapolation, the BOLD effects assessed by MRS on a spectral water peak could be compared to the changes seen in [32]. Moreover, upon depolarization induced by a stimulus, a Na^+^ gradient towards the intracellular milieu occurs. This analogy was used in the past with diffusion magnetic resonance imaging (dMRI) to verify that the dynamic swelling of neurons due to neuronal activation, induced a decrease of the apparent diffusion coefficient (ADC) [33]. Potential changes in physical environments inducing ADC changes would also affect T_2_. Since T_2_ changes also occur as a result of focal cortical activation, cell swelling or macromolecular changes may also be envisaged in the present study. Other groups also demonstrated important BOLD effects on the neuronal compartment, which origin was also attributed to cell swelling or macromolecular changes [7, 34]. At 9.4T, a negligible effect of intravascular BOLD was revealed [34]. If cell swelling due to neural activation is assumed, then vascular BOLD effects can also be neglected. Changes in water and NAA signals during optogenetic stimulation could therefore be related to an increase of the intracellular volume. Diffusion functional MRS (dfMRS) was also proposed as a new tool to separate the changes in properties of neuronal spaces from the hemodynamic response during neuronal activity [35]. Despite the potential of both diffusion techniques, many measurements remain to be performed to ensure their validation as probes for the direct measurement of neural activity. Besides, both methods require additional work to validate the underlying mechanisms and both are suspected to remain contaminated by residual BOLD signals. The present work participates in the effort to better identify genuine neural activity. A simple method was proposed to separate water and intracellular BOLD effects and will need to be validated.

### 4.3 In-vivo evaluation of BOLD effects

Evaluation of the *in-vivo* water and NAA peak amplitude changes as a function of time followed the stimulation paradigm during periods of photo-stimulation and periods of REST, mimicking BOLD changes usually seen in BOLD fMRI time courses. These findings suggest that estimating water and NAA peak height changes represents an acceptable means of identifying spectral BOLD effects in the rat cortex. On the contrary, changes in tCr peak amplitudes as a function of time did not follow the paradigm of stimulation. In a very recent study conducted at 9.4T in the human brain, tCr concentrations did not follow the visual stimulation paradigm, while Cr concentrations did [36]. These results confirm that BOLD effects may be counterbalanced by the conversion of PCr into Cr and vice versa as hinted by the absence of observable T_2_ changes in other studies [34]. In rodents, Medetomidine sedation may also be responsible for some modulation of metabolite levels [37].

*In-vivo,* stimulated spectra needed to be corrected for the BOLD effect without impacting the potential metabolite concentration changes due to optogenetic stimulation. Overall, the corrected amplitude of changes in metabolite concentrations upon stimulation as well as the direction (positive or negative) of these changes was in agreement with previous studies in rodents [13,14,15, 16, 26] for all the different methods. The best agreement with previous literature values on functional metabolic concentration changes was found for the line-matching procedure [13,14,15,26] with lb=0.5 Hz. Using the line-matching procedure with lb= 6 Hz, changes in Asp and Glu concentrations were above −1 μmol/g (Table 3), which were largely overestimated compared to previous reports [13,14,15]. Moreover, the linewidth-matching correction with lb=6 Hz demonstrated to be erroneous (**Fig. 7**). Given these results, the line-matching procedure using lb=0.5 Hz appears the most suitable for correcting BOLD effects without affecting metabolic concentration changes due to optogenetic stimulation in the present study.

### 4.4 BOLD-free difference spectra

Here, findings with BOLD free difference spectra were considered to remain only indicative of the polarity of changes due to low SNR. Nevertheless, direct LCModel fitting led to reasonable quantitative estimates for Glu and Asp concentration changes (Table 4) compared to changes reported in Table 3. The CRLB for Lac were 40 % and thus were not reported.. Observations were qualitatively performed by visual inspection of the overlaid linewidth-corrected (lb= 0.5 Hz) stimulated and REST spectra. Subtraction of the resting spectrum from the spectrum acquired during photo-stimulation and corrected for BOLD effects resulted in small positive and negative peaks representing stimulus-related changes in several metabolites. Positive (Glu, GSH, Lac) and negative (Asp, Lac) peaks representing metabolite concentration changes were identified at specific chemical shift positions confirming quantitative findings using LCModel fittings reported in Table 3.

## 5. CONCLUSION

In conclusion, BOLD effects can be quantified using ^1^H-fMRS in the rat cortex using ^1^H-fMRS. Linewidth-matching techniques remain the most reliable correction for eliminating false positive errors in metabolite concentration estimates provided the extent of T_2_*-induced effects is characterized for the precise determination of the line broadening factor. The ensuing calculation of a BOLD-corrected difference spectrum should then be more straightforward and more specific. Alternatively, BOLD-related spectral changes may be useful to further identify differences between cortical extravascular compartments upon activation. As pointed out recently (38), novel T_2_* methods are needed to better correct for BOLD effects in ^1^H-fMRS. At the same time, a better characterization of BOLD effects independent of neurovascular coupling may be useful for an improved assessment of direct neuronal activity.

## 6. Acknowlegements

The author would like to thank the Translational Research Institute Center for providing animals. All fMRS Matlab routines were adapted from previous routines developed in the Centre d’Imagerie Biomédicale (Lausanne, Switzerland) by the first author.

